# A unified multiscale modelling framework to explore the brain excitatory-inhibitory balance: application to multiple sclerosis

**DOI:** 10.64898/2025.12.03.692031

**Authors:** Gökçe Korkmaz, Roberta Maria Lorenzi, Francesca Ravera, Adnan Alahmadi, Anita Monteverdi, Baris Kanber, Ferran Prados Carrasco, Egidio D’Angelo, Fulvia Palesi, Ahmed Toosy, Claudia A.M. Gandini Wheeler-Kingshott

**Author notes:** **Corresponding Author:** Claudia Gandini Wheeler-Kingshott.

## Abstract

Balanced excitation and inhibition are essential for brain dynamics, and their disruption can lead to network dysfunction in neurological diseases. Here, we present a conceptually unified multiscale brain modelling framework combining Dynamic Causal Modelling (DCM) applied to task and resting-state functional Magnetic Resonance Imaging (fMRI) data and The Virtual Brain (TVB) to characterise the excitatory/inhibitory balance of the brain. We applied the framework to a visuomotor brain subnetwork in a cohort of 9 healthy controls and 17 people with multiple sclerosis (pwMS). Acquired data included an event-related task fMRI experiment with variable grip force, resting-state fMRI, and diffusion-weighted imaging. The visuomotor network comprised the bilateral primary visual cortex (V1), left primary motor cortex (M1), supplementary motor and premotor cortex (SMAPMC), cingulate cortex (CC), superior parietal lobule (SPL), and right cerebellar lobule VI (CR). Results from DCM showed that while the overall network architecture was preserved in MS, there were significant alterations in the excitatory/inhibitory nature of effective connectivity: at rest, a statistical change was observed in CR-to-V1 connectivity, which was inhibitory in healthy volunteers but excitatory in MS. During task, effective connectivity feedback, including cerebellar self-connection, was positive in healthy volunteers but negative in MS and became increasingly dysregulated with higher motor demand. Alterations in functional and effective connectivity were associated with behavioural performance (task reaction time) and clinical measures (disability severity). At the overall group level, TVB parameters linked reduced NMDA-mediated excitatory gain to slower task responses. Moreover, integrating DCM and TVB demonstrated that higher global excitatory gain was associated with stronger task-engaged effective connectivity across sensorimotor and visuomotor pathways, linking network-level excitability captured by TVB to context-dependent reconfiguration of directed interactions and to connection-level strength revealed by DCM.

## 1. Introduction

Balanced excitation and inhibition is a fundamental property of neuronal systems that supports stable activity across spatial scales. Disruptions of this balance can emerge locally, yet it can impact large-scale brain dynamics, influencing directed interactions between cortical and subcortical regions. Recent advances in computational neuroscience enable the integration of processes across micro-, meso-, and macroscale levels using complementary modelling approaches (De Schepper et al., 2022; Geminiani et al., 2018). At the mesoscale, population dynamics are described using models that explicitly incorporate excitatory and inhibitory interactions (Geminiani et al., 2024; Kennedy et al., 2024), whereas at the macroscale, whole-brain activity inferred from functional Magnetic Resonance Imaging (fMRI) is typically modelled using neural mass or mean-field formulations constrained by region to region structural connectivity (Deco et al., 2008; Kennedy et al., 2024). Within this framework, the sign and strength of interregional interactions provide a parsimonious way of characterising network dynamics, but remain difficult to infer consistently across experimental contexts.

Dynamic Causal Modelling (DCM) offers a principled generative framework for estimating effective connectivity, i.e. directed interactions between brain regions, from fMRI data (Friston et al., 2003a). In parallel, The Virtual Brain (TVB) provides a platform to simulate whole-brain dynamics by combining structural connectivity with neural mass models parameterised in terms of excitatory and inhibitory synaptic gains (Ritter et al., 2013; Sanz-Leon et al., 2013). While both approaches have been widely used, they are typically applied in isolation: DCM focuses on pairwise interactions estimated from observed data, whereas TVB emphasises global network behaviour emerging from generative simulations. The relationship between these two levels of description remains insufficiently characterised, particularly with respect to how local directed interactions relate to macroscopic network regimes.

Multiple Sclerosis (MS) provides a relevant testbed to address this question. MS is a chronic neuroinflammatory disease associated with widespread structural damage and functional reorganisation (Filippi et al., 2019; Lassmann et al., 2012; Ricciardi et al., 2023; Rocca & Filippi, 2007). fMRI studies have consistently demonstrated altered network organisation during both rest and task (Alahmadi et al., 2021; Alharthi & Almurdi, 2023; Mezzapesa et al., 2008; Rocca et al., 2012), including nonlinear modulation of visuomotor responses with increasing motor demand (Alahmadi et al., 2017). However, these findings are largely descriptive and do not provide direct insights into the underlying mechanisms governing interregional interactions. In particular, it remains unclear whether disease-related changes reflect differences in the magnitude of coupling, changes in its directionality, or shifts in the inferred excitatory/inhibitory balance.

Computational approaches offer a means to address these limitations by estimating latent neuronal processes from measured signals (D’Angelo & Jirsa, 2022; Frässle et al., 2018). DCM studies in MS have reported alterations in effective connectivity, often with substantial inter-subject variability (Au Duong et al., 2005; Mansoory et al., 2022; Savini et al., 2019a). In parallel, a small number of studies have used TVB to infer global changes in network dynamics (Martí-Juan et al., 2023; Sorrentino et al., 2024). However, these approaches have not yet been combined to examine how connection-level estimates of effective connectivity relate to network-level parameters within a conceptually unified modelling framework, nor how these relationships depend on behavioural context (e.g. task vs rest).

In this study, we address this gap by integrating DCM, spectral DCM (sp-DCM), and TVB within a single modelling framework applied to a visuomotor network in people with MS (pwMS) and healthy volunteers (HV). We focus on three questions: (i) whether the inferred sign (excitatory vs inhibitory) of effective connectivity differs between groups; (ii) whether these differences depend on context (task vs resting-state); and (iii) how connection-level estimates relate to global model parameters and behavioural or clinical measures. To this end, we analyse task-based fMRI, resting-state fMRI, and diffusion MRI from a previously acquired dataset (Alahmadi et al., 2021), focusing on a visuomotor network including bilateral primary visual cortex (V1), left primary motor cortex (M1), supplementary motor and premotor cortex (SMAPMC), cingulate cortex (CC), superior parietal lobule (SPL), and right cerebellar lobule VI (CR) (Lorenzi et al., 2025). DCM and sp-DCM are used to estimate effective connectivity under task and resting conditions, respectively, while TVB is employed to model large-scale dynamics constrained by structural connectivity. The resulting parameters are then related to behavioural performance and clinical disability using Parametric Empirical Bayes (PEB) (Zeidman et al., 2019) and regression model approaches.

## 2. Methods

This section describes the unified brain modelling framework applied to both HV and pwMS. As shown in Figure 1, all participants underwent functional and diffusion MRI preprocessing, followed by complementary DCM and TVB analyses detailed below.

**Figure 1.**
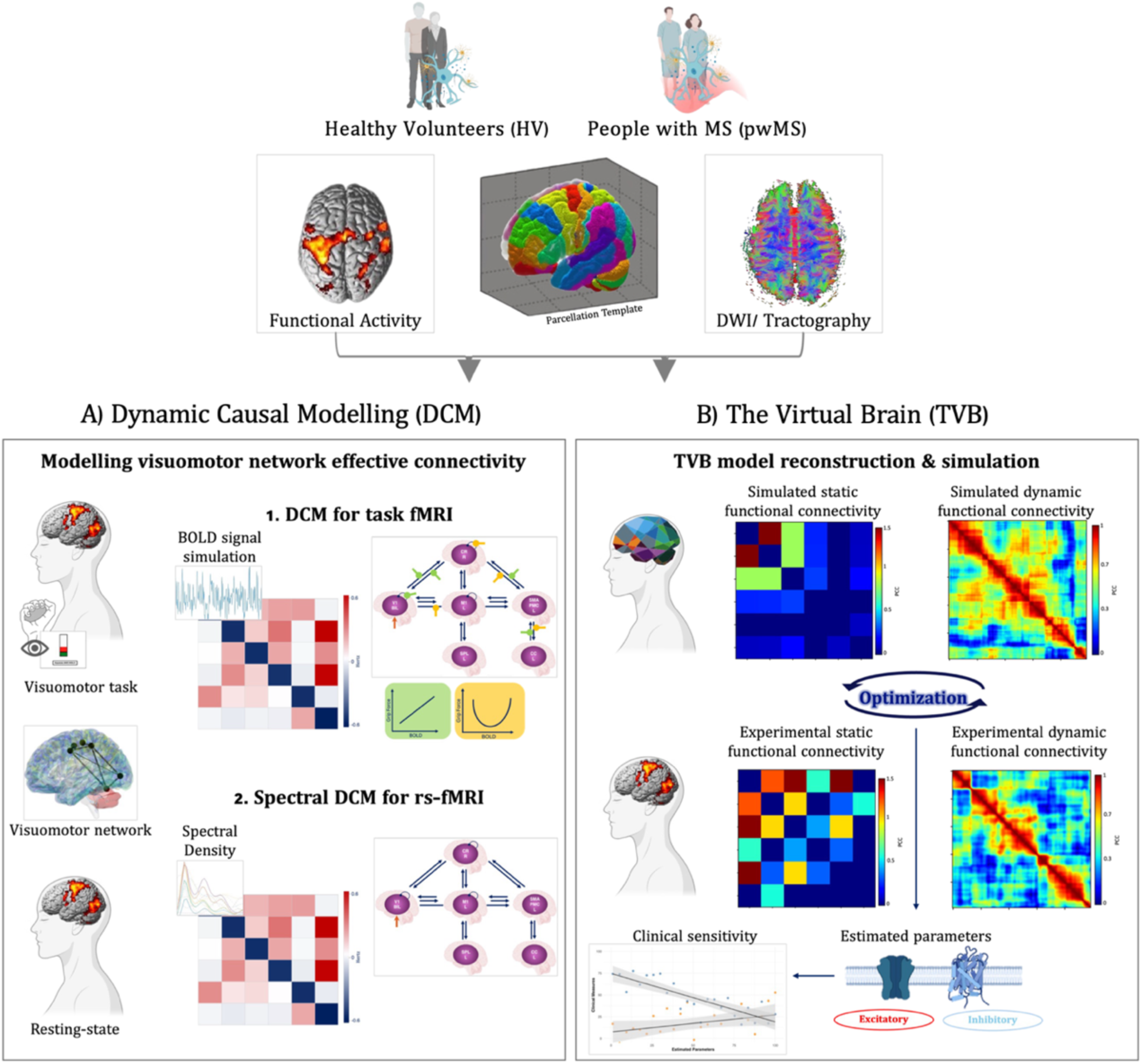
Conceptually unified brain modelling framework. The brain modelling approach included pipelines for Dynamic Causal Modelling (DCM) (A) and The Virtual Brain (TVB) (B) analyses using combinations of structural diffusion-weighted imaging (DWI), task-based and resting-state fMRI data on people with multiple sclerosis (pwMS) and healthy volunteers (HV). The same visuomotor network, with grey matter nodes parcellated using a Brodmann/SUIT ad hoc atlas, is used in all analyses. DCM for task fMRI (A.1) computes the effective connectivity dynamics of the defined visuomotor network and its modulation with different grip force levels using task fMRI, while spectral DCM (sp-DCM) is applied to resting-state fMRI (A.2) to compute the effective connectivity in the same network based on frequency fluctuations of resting-state fMRI. TVB simulates subject-specific network functional dynamics starting from structural connectivity and optimizing the model of neuronal function associated to the network nodes against the experimental functional connectivity (both static and dynamic). Clinical (EDSS) and behavioural (reaction times) parameters are regressed against model parameters. Statistically significant TVB parameters will be explained in terms of pairwise effective connectivity from DCM and sp-DCM.

### Participants

Participants were selected from a cohort recruited as part of a previous study (Alahmadi et al., 2021): nine right-handed HV [4 females, 5 males; age = 37 ± 6 years], selected based on availability of resting-state fMRI, DWI, and task fMRI data; and all fourteen right-handed pwMS (Relapsing-Remitting MS, RRMS), with three additional participants for a total of 17 MS subjects [12 females, 5 males; age = 35 ± 7 years; median (range) EDSS = 3.5 (1.5-6.5)]. For the TVB analyses, a subset of these participants was used (7 HV and 14 pwMS), based on the availability of the data necessary for model construction. Participants’ handedness was evaluated using the Edinburgh handedness inventory questionnaire (Oldfield, 1971). All subjects gave informed consent, and the study was approved by the local research and ethics committee and was performed in line with the principles of the Declaration of Helsinki.

### MRI Acquisition & fMRI Study Design

MRI data were acquired for all participants using a 3 Tesla Philips Achieva scanner (Philips Healthcare, Best, The Netherlands) equipped with a 32-channel receive-only head coil. All image volumes were aligned to the bicallosal line. The imaging protocol included structural, functional (task-based and resting-state), and DWI acquisitions. Details are reported in Supplementary Table 1.

Task-based fMRI data were collected using a BOLD-sensitive T2*-weighted echo-planar imaging sequence while participants performed an event-related power grip task (TR/TE = 2500/35 ms, flip angle = 90°, voxel size = 3 × 3 × 2.7 mm³). Participants used their right (dominant) hand to squeeze an fMRI-compatible rubber ball in response to a visual cue. The task consisted of 75 active trials distributed evenly and randomly across five GF levels (20%, 30%, 40%, 50%, and 60% of each participant’s maximum voluntary contraction). Actual task execution timings were recorded for each subject, and reaction times (delays between cue onset and motor execution) were calculated and z-score transformed for subsequent analyses.

Resting-state fMRI data were acquired using the same imaging parameters as the task-based fMRI sequence, with a reduced number of volumes. Structural imaging included high-resolution T1-weighted and PD/T2-weighted sequences for anatomical characterization and lesion detection.

DWI was performed using a high angular resolution diffusion imaging protocol. Axial-oblique DWI data were acquired with a cardiac-gated spin-echo (SE) EPI sequence (TR ≈ 24,000 ms, TE = 68 ms, flip angle = 90°, voxel size = 2 × 2 × 2 mm³).

### Data preprocessing of raw MRI data

#### White matter lesions

White matter hyperintensity were delineated by hand on the PD/T2-weighted scans for all MS participants, using the JIM software version 9 *(Xinapse Systems)* according to best practice.

#### Structural scans

Structural (3D T1-weighted) images were first lesion filled using the lesion masks according to (Prados et al., 2016). They were then used for the fMRI and DWI processing as described below.

#### fMRI data

fMRIPrep 23.2.1, a containerized standardized pipeline based on Nipype (Esteban et al., 2019), was used for both task and resting-state data. Detailed steps are reported in the supplementary materials.

#### DWI data

Preprocessing of the single-shell DWI data was conducted following in-house procedures, e.g. described by (Klein et al., 2012): the seven b=0 volumes were averaged to create a mean b=0 volume, which was used as a reference to which register each of the 61 DW volumes, using NiftyReg (http://niftireg.sf.net). EPI distortions were corrected using Brain Suite (Charalambous et al., 2019) while motion correction was performed with eddy (Andersson & Sotiropoulos, 2016). DWI data were used to reconstruct the subject-specific structural connectomes: a whole-brain tractogram was obtained with an anatomically constrained tractography (Smith et al., 2012) (MRtrix3, http://www.brain.org.au/software/mrtrix/) with 30 million streamlines, the 5-tissue-type (5TT) segmentation and a dynamic seeding approach. The registration between 3D T1 images and DWI data was performed following the procedure described by (Klein et al., 2012). First, 3D T1 images were non-linearly registered to Montreal Neurological Institute (MNI) space. Tissue segmentation (e.g., grey matter, white matter, and cerebrospinal fluid) was then performed in the native 3D T1 space to preserve anatomical accuracy. For diffusion processing, the 3D T1 images were affine registered to a pseudo-T1 image derived from the corresponding PD/T2-weighted scans. Subsequently, the T2-weighted images were linearly and then non-linearly registered to the mean b=0 image. All transformations were concatenated to allow mapping between native 3D T1 space, DWI space, and MNI space in both directions. All registrations were performed using the NiftyReg.

### Network definition & time-series extraction

#### For all modelling pipelines

A 6 grey matter regions (nodes) atlas of the visuomotor network was defined based on previous publications reporting results on the same visuomotor fMRI task used in this cohort (Alahmadi et al., 2016; Lorenzi et al., 2025). The visuomotor network nodes included bilateral V1, left M1, SMAPMC, CC, SPL, and right CR. Regional masks were derived from Brodmann area parcellations (Mai & Milan, 2017) and SUIT cerebellar atlas (Diedrichsen, 2006) (Figure 2, DCM, panel A and TVB panel A). This network atlas was used consistently in both DCM, sp-DCM and TVB analyses.

**Figure 2.**
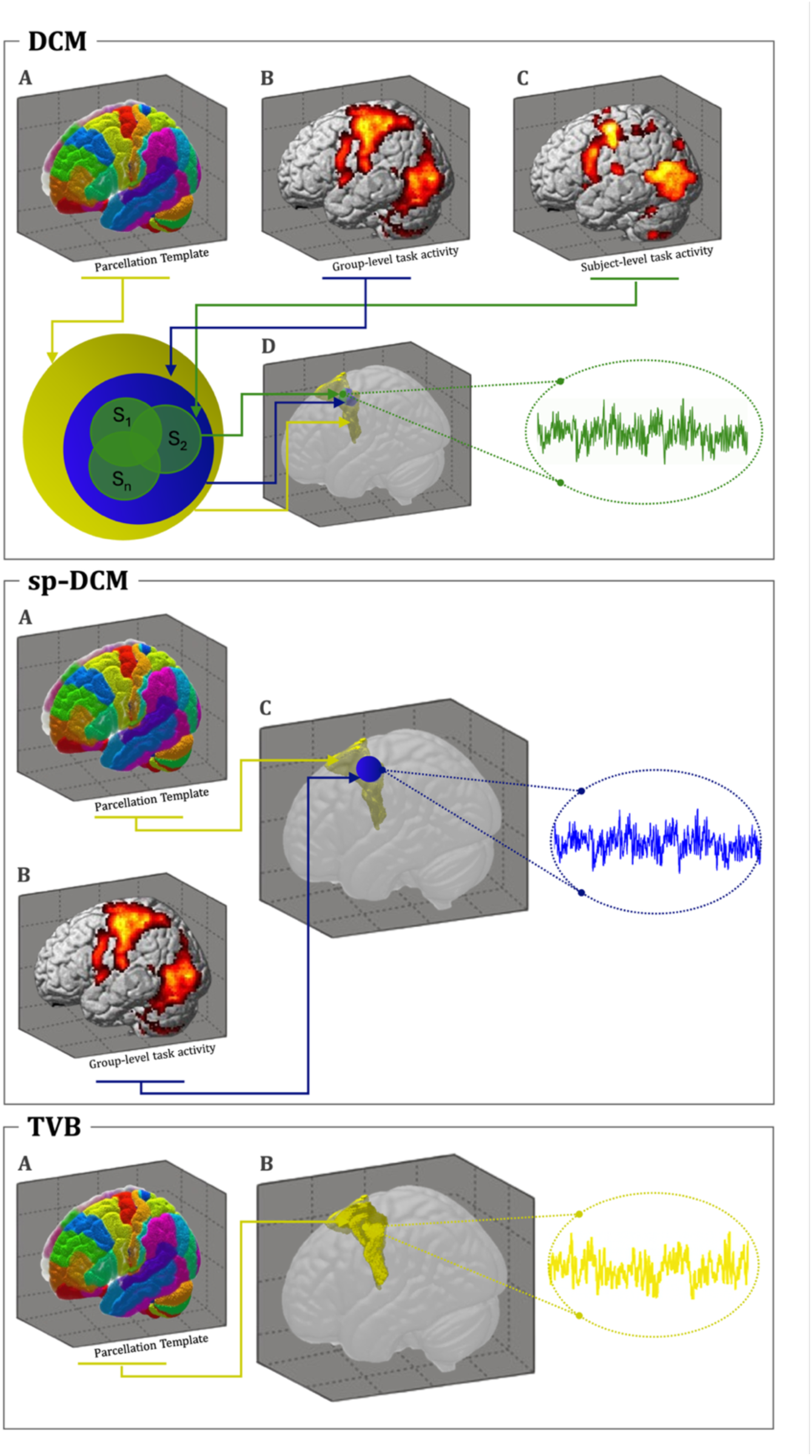
fMRI Time-Series Extraction Pipeline. DCM panel. For each node of the visuomotor network, the pipeline included subsequently restricting constraints: **(A)** anatomical constraints defined using an ad hoc atlas (Brodmann/SUIT), **(B)** functional constraints defined as spheres of 15 mm radius centred on the peak of the task-based group-level activity for both HV and pwMS separately, **(C)** geometrical constraints defined as the subject-specific peak activity in a sphere of 6 mm radius, within the intersection of A and B. **(D)** Example of 3D visualization of the anatomical (yellow), functional (blue) and geometrical (green) constraints for the left supplementary motor area & premotor cortex (SMAPMC). The time-series in the left circle is extracted from the final volume of interest (VOI) (illustrated in green). **sp-DCM panel -** The identified coordinates with the DCM pipeline, using task fMRI from HV only, at the group level, were applied to define the VOI for resting-state fMRI signal extraction. **TVB panel** - an ad hoc atlas masks were used to define VOI for time-series extraction for TVB.

#### DCM analysis

Task-based group-level activation masks were identified through second-level analysis of HV and pwMS separately. Peak coordinates within each region, and a sphere of 15 mm radius centred on the group level peak was intersected with the corresponding atlas mask (Figure 2, DCM, panel B; Table 1). Subject-specific peak activity was then identified within the group-level mask for each participant, and 6 mm radius sphere centred on the individual peak was intersected with the group mask (Figure 2, DCM, panel C). Finally, the regional time-series was computed only from those voxels belonging to the conjunction of the three masks from these steps, defining the final Volume of Interest (VOI) (Figure 2, DCM, panel D).

**Table 1.** Volume of Interest (VOI) details. Threshold indicates the uncorrected significance level used on task fMRI results to select activated voxels included in the VOI. The coordinates for task fMRI are defined at group-level separately for HV and pwMS. All VOIs were defined as spheres with a radius of 6 mm. The coordinates for resting-state fMRI are identified starting from those of HV in task fMRI.

| Region | Threshold task | Coordinates task fMRI (HV) | Coordinates task fMRI (pwMS) |
| --- | --- | --- | --- |
| Primary motor cortex (M1), left | 0.001 | [-40; -21; 54] | [-38; -18; 54] |
| Supplementary motor area & premotor cortex (SMAPMC), left | 0.001 | [-4; -6; 54] | [-3; -6; 54] |
| Superior parietal lobule (SPL), left | 0.001 | [-37; -36; 45] | [-35; -37; 44] |
| Cingulate cortex (CC), left | 0.001 | [-7; -3; 42] | [-4; -1; 40] |
| Primary visual cortex (V1), bilateral | 0.01 | [14; -54; -21] | [-2; -83; 3] |
| Cerebellar lobule VI (CR), right | 0.001 | [23; -99; 0] | [16; -56; -19] |

#### sp-DCM analysis

VOI coordinates obtained from the analysis of task fMRI of HV (Figure 2, sp-DCM, panel A) were used to define the centre of a 6 mm sphere used to extract time-series from resting-state fMRI data (Figure 2, sp-DCM, panel B) in the atlas regions.

#### TVB analysis

TVB requires as input two functional connectivity matrices from resting-state fMRI data, i.e. a static (FC) and a dynamic functional connectivity (FCD) matrix, and two structural connectivity (SC) matrices from tractography, i.e. one weighted on streamline count and the other on streamline length. First, the time-series were extracted from the visuomotor nodes (Figure 2, TVB) and, the Pearson Correlation Coefficient (PCC) was calculated between pairs of them and transformed as Fisher’s z-values, thresholded at 0.1206 to retain only significant correlations (Palesi et al., 2020). This static experimental FC reflects the overall level of connectivity between pairs of brain regions during the resting-state condition. Experimental FCD, which represents the dynamic nature of FC, was obtained correlating across time several FCs computed over a sliding window of 40 seconds. This window was incrementally shifted by 1 TR, resulting in a series of 105 experimental FC windows. The two SC matrices were created by extracting streamlines connecting pairs of regions (i.e., nodes) of the visuomotor network atlas from the whole-brain tractogram. Only the contralateral cerebro-cerebellar tracts between cerebellar lobule VI and the other cerebral regions were considered and extracted following the anatomically-correct procedure outlined in (Palesi et al., 2015, 2017).

### Brain Modelling

The brain modelling approach (Figure 1) comprises three pipelines: DCM, sp-DCM, and TVB, integrating structural, functional, and effective connectivity. Analyses were performed in HV and pwMS, with correlations between model parameters and clinical disability also examined. Details follow.

#### Dynamic Causal Modelling

DCM analyses task fMRI data to model effective connectivity between pairs of regions (Zeidman, Jafarian, Corbin, et al., 2019). V1 was set as the driving input according to the nature of the task and our previous work (Lorenzi et al., 2025). DCM12.5 was used as implemented in SPM12 (MATLAB 2022b).

Predicted neural dynamics were transformed into BOLD signals using the haemodynamic forward model (Friston et al., 2000; Stephan et al., 2007b), to extract endogenous coupling between pairs of regions, independently of the context (i.e., fixed (intrinsic) effective connectivity). After, the impact of the experimental stimuli (i.e. GF levels) was modelled as a 1^st^ order (linear) or 2^nd^ order (non-linear) modulation on the strength of the fixed effective connectivity between regions. Fixed effective connectivity was examined as the first step (S1) and modulations on fixed effective connectivity as second step (S2). For both S1 and S2, the pipeline described by (Lorenzi et al., 2025) for action execution during the GF task was followed, separately for HV and pwMS. In S1, the same previously published network architectures (Supplementary Figure 1) were tested. Bayesian model selection (BMS) was used to select the best-fitting model for a different set of hypotheses at a group-level (Penny et al., 2010). After identifying the best-fitting model for both HV and pwMS (S1), each model configuration was further investigated by estimating the posterior probability (Pp) of each connection (S1) and modulation (S2) specified in the model architecture. In S2 both 1^st^ and 2^nd^ order modulations were imposed on each pair of connections on the winning S1 model, focusing on the visuo-to-plan loop (V1-CR-SMAPMC-CC); different combinations were tested following a stack procedure that selected the most probable modulation for a certain connection and moved along the loop (Supplementary Figure 2). The overall process quantified the model likelihood, assessed as model evidence using the experimental fMRI time-series data. Then, Random Effects-BMS (RFX-BMS) was employed to determine whether the winning model varied across subjects (Penny et al., 2010). Bayesian Model Averaging (BMA) was employed to integrate the statistical structure by computing a weighted average of each connectivity strength across the model space (Friston et al., 2016).

#### Spectral Dynamic Causal Modelling

sp-DCM is a DCM variant for resting-state fMRI that fits the cross-spectral density (frequency-domain representation of BOLD signals) to estimate interregional coupling. By modelling these frequency components, it captures how spontaneous neural fluctuations give rise to intrinsic connectivity without external inputs. (Li et al., 2011; Novelli et al., 2024). sp-DCM (DCM12.5 implemented in SPM12) was applied on the same winning model architecture from DCM (S1), for both HV and pwMS to capture frequency-specific dynamics in the same visuomotor network. BMA was then applied across subjects to estimate effective connectivity strengths at rest.

#### The Virtual Brain Modelling

The TVB (version 2.7.2) workflow, as reported in (Monteverdi et al., 2022), was followed (Figure 1.B). The Wong-Wang neural mass model (Deco et al., 2014) was chosen to characterize local cortical activity through two coupled populations representing excitatory (glutamatergic) and inhibitory (GABAergic) neurons. This model is characterized by four parameters: i) the global coupling *(G)* that helps quantifying the overall integration capacity of the brain and determines how strongly brain regions influence each other; ii) the NMDA synaptic weight (J_NMDA_) that determines the strength of the excitatory (glutamatergic) synaptic connections; iii) the inhibitory coupling parameter (J_i_) that controls the strength of inhibitory (GABAergic) synaptic connections; and iv) the recurrent excitatory strength of each brain region (w_+_) that shows how strongly excitatory populations excite themselves.

The SC was used to connect all the nodes of the visuomotor network and each node was associated with a Wong-Wang model (Wong et al., 2006). Then, TVB generated a simulated FC and FCD for different combinations of model parameters (*G*, J_NMDA_, J_i_, w_+_) that were optimized using an iterative fitting approach described in (Good et al., 2022): first the PCC between simulated and experimental FC was maximized, then Kolmogorov-Smirnov (KS) distance between simulated and experimental FCD was minimized. This optimisation was run for all participants to the study, obtaining a set of subject-specific model parameters for the network.

### Statistical Analysis and Disease Sensitivity

An independent sample t-test was conducted to compare the best-fitting S1 model effective connectivity matrices between the HV and pwMS for both DCM and sp-DCM using R (version 2024.12.0). P-values were corrected for multiple comparisons using the false discovery rate (FDR) and the significance threshold was set at 0.05.

PEB framework was used for group-level DCM analysis with clinical/behavioural scores (Friston et al., 2015; Friston et al., 2016). While DCM provides valuable estimates of subject-specific effective connectivity that can differ between groups, understanding the clinical relevance of such differences requires an additional step that can be achieved by applying PEB analysis; providing a hierarchical Bayesian approach that accounts for between-subjects’ effects, identifies connectivity alterations across individuals and characterises how experimental or clinical variables are associated with effective connectivity differences. All statistical analysis assumed that the reaction time measured from the GF task performance can be considered a behavioural test of functional integrity for all subjects, while EDSS was investigated considering only pwMS. PEB was applied across all subjects: (i) to investigate group-level effects of the GF task performance, assessed through the z-score transformed reaction time, and (ii) only to pwMS for EDSS to examine the association of disease severity with the effective connectivity between specific nodes. After, the same analysis was repeated for sp-DCM.

Regarding TVB parameters, the Shapiro-Wilk test was applied to assess the normality of continuous variables, including age, sex, and all TVB-derived parameters. To identify the most relevant predictors of clinical disability (EDSS) and the reaction time, bidirectional stepwise regressions were run in R (version 2024.12.0), using the Akaike Information Criterion (AIC) as the selection criterion. Bidirectional stepwise procedures were performed independently to ensure the robustness of the predictors’ selection. A p-value threshold of 0.05 was used to determine statistical significance for a variable inclusion in the final model. A *post-hoc* analysis used PEB to examine which pairwise effective connectivity values were explained by the network-level TVB parameter associated with behaviour. PEB was run using both task and resting-state data.

### Reproducibility and data accessibility

The analyses pipelines used for this study are available on GitHub/GokceKo/DCM-TVB-MS. The data supporting this study are available from the corresponding author upon reasonable request and subject to data transfer agreements. Data are stored in a local controlled-access repository.

## 3. Results

We observed complex alterations in effective connectivity within the visuomotor network in MS, predominantly involving cerebellar-cortical interactions. While TVB simulations indicated no marked changes in global network-level parameters, these models captured clinically and behaviourally relevant associations. Effective connectivity analyses further revealed context-dependent differences between HV and pwMS, with stronger excitatory influences during task performance in HV and a relatively more inhibitory network profile at rest in pwMS. In summary, we found that:

- DCM supports the same effective connectivity network architecture in HV and pwMS (S1.4, Figure 3A).
- Compared with HVs, pwMS shows:

○ changes in the excitatory and inhibitory nature of effective connectivity between task and rest (Figure 3B), with connection-specific patterns that differ across context (Figure 3C). In particular, the CR-to-V1 effective connectivity is significantly different between HV and MS during rest, with an inhibitory nature in HV and excitatory in MS;
○ alterations in the GF-BOLD modulation of the effective connectivity between pairs of regions (Figure 4), with the feedback from CC to V1 via SMAPMC and CR being negative in pwMS, including the CR self-connection, while positive in HV.
- For clinical outcomes:

○ effective connectivity between pairs of regions (e.g. V1 to M1, V1 to CR, M1 to SMAPMC) is associated with EDSS and reaction time;
○ TVB excitatory synaptic gain (J_NMDA_) is inversely correlated with the reaction time (Figure 5).
- A post-hoc analysis revealed that pairwise effective connectivity values were explained differently by network-level J_NMDA_ depending on the task or resting-state context (Figure 6).

**Figure 3.**
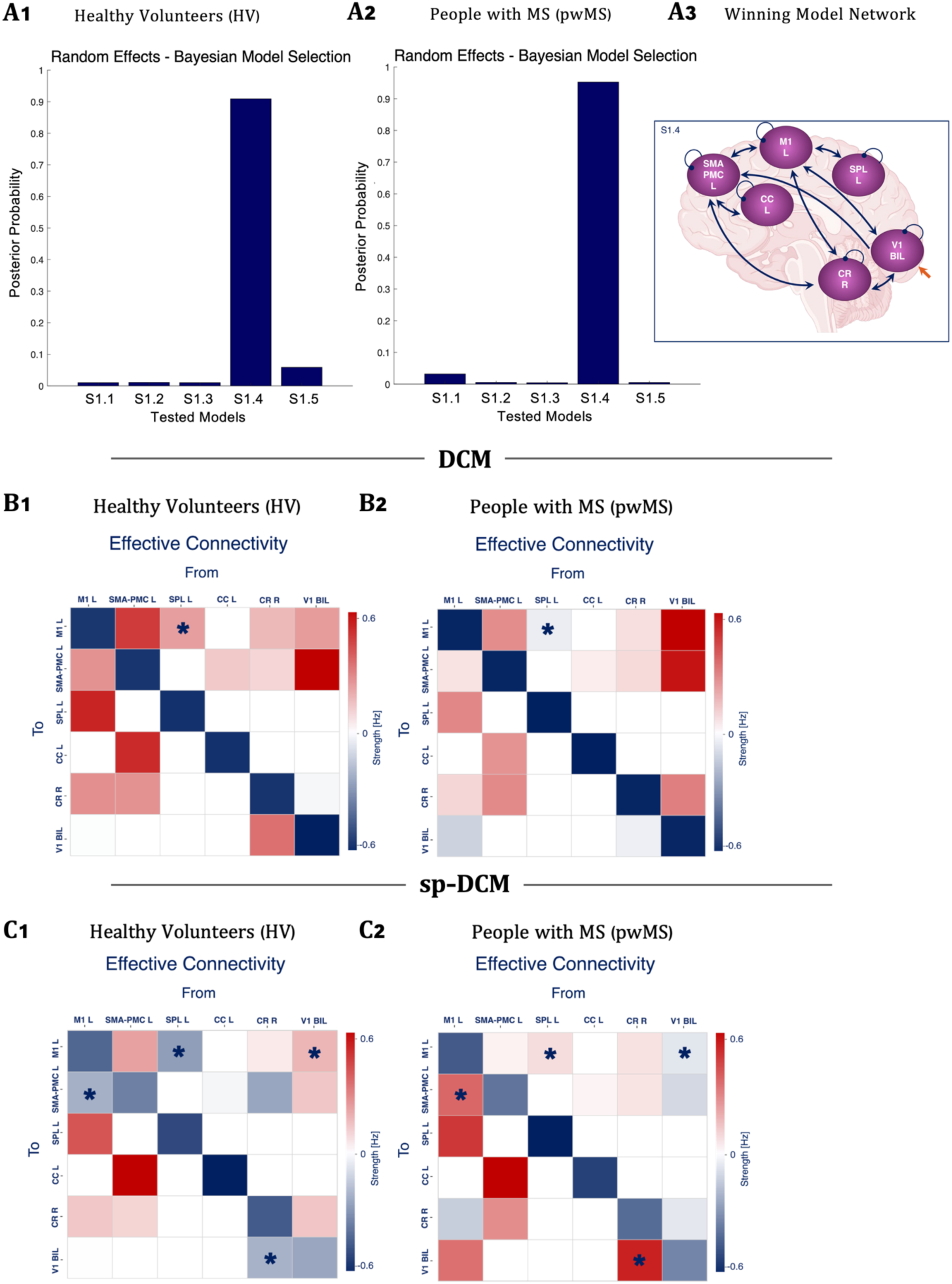
DCM and sp-DCM results on 0ixed effective connectivity (S1). **A)** Bayesian Model Selection results for HV (left panel) and pwMS (middle panel), with visual representation of the winning model network architecture (right panel). Model S1.4 is the best-fitting model for DCM with a posterior probability > 90% for both HV and pwMS. **B)** DCM fixed effective connectivity results for the winning model (S1.4) in Hz. **C)** sp-DCM fixed effective connectivity results for S1.4 in Hz. The red colour indicates excitation whereas the blue colour indicates inhibition. White values indicate connections that are switched off in the corresponding network. Self-connections are unitless log-scaling parameters that multiply a default value of −0.5 Hz. Asterisk * indicates statistically significant differences between groups.

**Figure 4.**
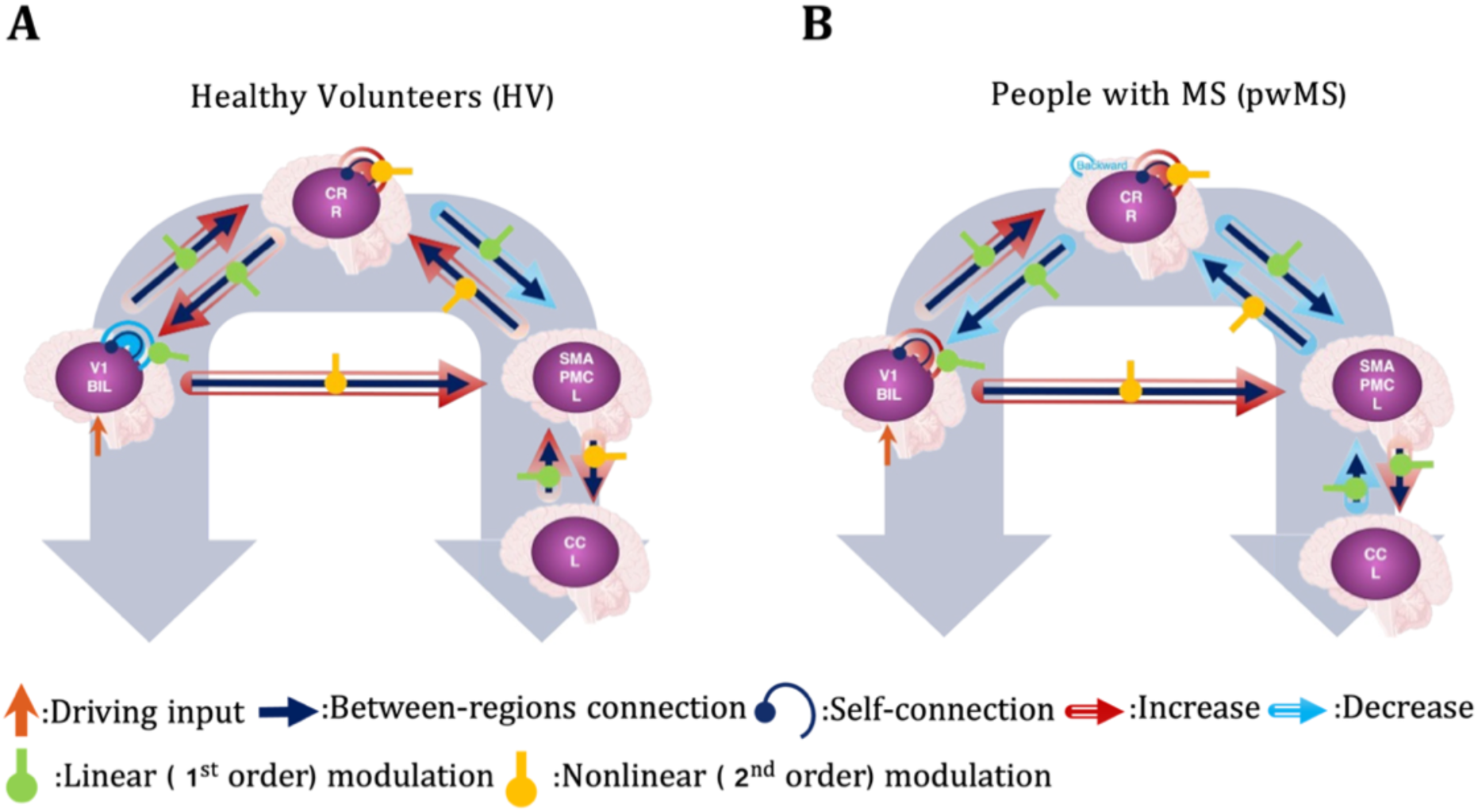
DCM results for modulations on 0ixed effective connectivity (S2). Modulations on the fixed effective connectivity for HV (A) and pwMS (B) are shown for both the forward (from V1 to CR to SMAPMC to CC) and the backward (from CC to SMAPMC to CR to V1) pathways. **For HV (panel A):** In the forward pathway, only from CR to SMAPMC showed a negative modulation by the GF-BOLD relationship; while all others showed positive modulation. Effective connectivity exhibited a linear modulation with grip force from V1 to CR, while the CR self-connection showed a nonlinear modulation. Connectivity from CR to SMAPMC also varied linearly with grip force, whereas connectivity from SMAPMC to CC and from V1 to SMAPMC demonstrated nonlinear modulation. In the backward pathway, all modulations were positive, apart from the V1 self-connection, indicating that an increasing GF-BOLD relationship generally enhanced the strength of the effective connectivity between pairs of regions. The effective connectivity modulation from SMAPMC to CR and within the CR self-connection was nonlinear, while the remaining connections showed linear responses. **For pwMS (panel B):** In the forward pathway, only the modulation from CR to SMAPMC was negative with respect to GF-BOLD relation; all others showed positive responses. Effective connectivity varied linearly with grip force from V1 to CR, became nonlinear in the CR self-connection, and returned to linear modulation from CR to SMAPMC and from SMAPMC to CC. Nonlinear modulation was also present from V1 to SMAPMC. In the backward pathway, modulations were negative, indicating reduced effective connectivity with respect to increasing grip force-BOLD relation except for the V1 self-connection. Nonlinear modulations appeared from SMAPMC to CR and in the CR self-connection, whereas other connections remained linear. *V1 = bilateral primary visual cortex, M1 = left primary motor cortex, SMAPMC = left supplementary motor area and premotor cortex, CC = left cingulate cortex, SPL = left superior parietal lobule, CR = right cerebellum*.

**Figure 5.**
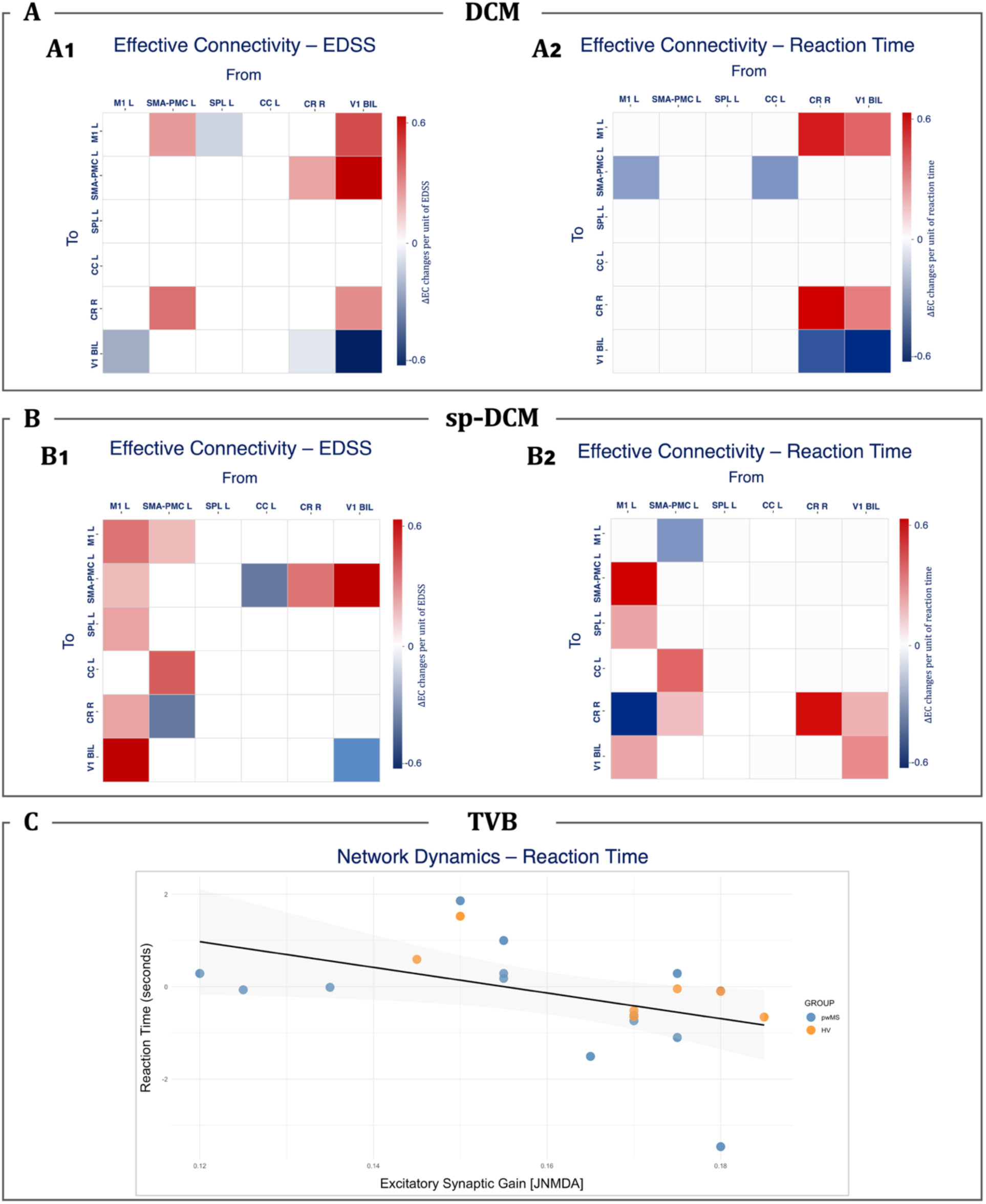
Network dynamics associated with EDSS and reaction time. Effective connectivity estimated on the DCM winning model (S1.4). Pairs of regions associated with EDSS (A1, B1) and reaction time (A2, B2) are identified by PEB analysis with a posterior probability greater than 0.95. Red indicates the increase in effective connectivity per unit change of scores, while blue indicates the decrease in effective connectivity per unit change of scores. V1 = bilateral primary visual cortex, M1 = left primary motor cortex, SMAPMC = left supplementary motor area and premotor cortex, CC = left cingulate cortex, SPL = left superior parietal lobule, CR = right cerebellum. TVB panel shows a significant negative association between J_NMDA_ and the reaction time calculated as the delay in squeezing the ball for all participants as identified by a bidirectional stepwise linear regression model with all TVB parameters plus age and sex as independent variables (p = 0.034, R² = 0.216). Each point represents one participant. Orange dots are the HV scores, and blue dots are pwMS. The modelled regression line (in black) and 95% confidence interval (grey shadow) are shown (C).

**Figure 6.**
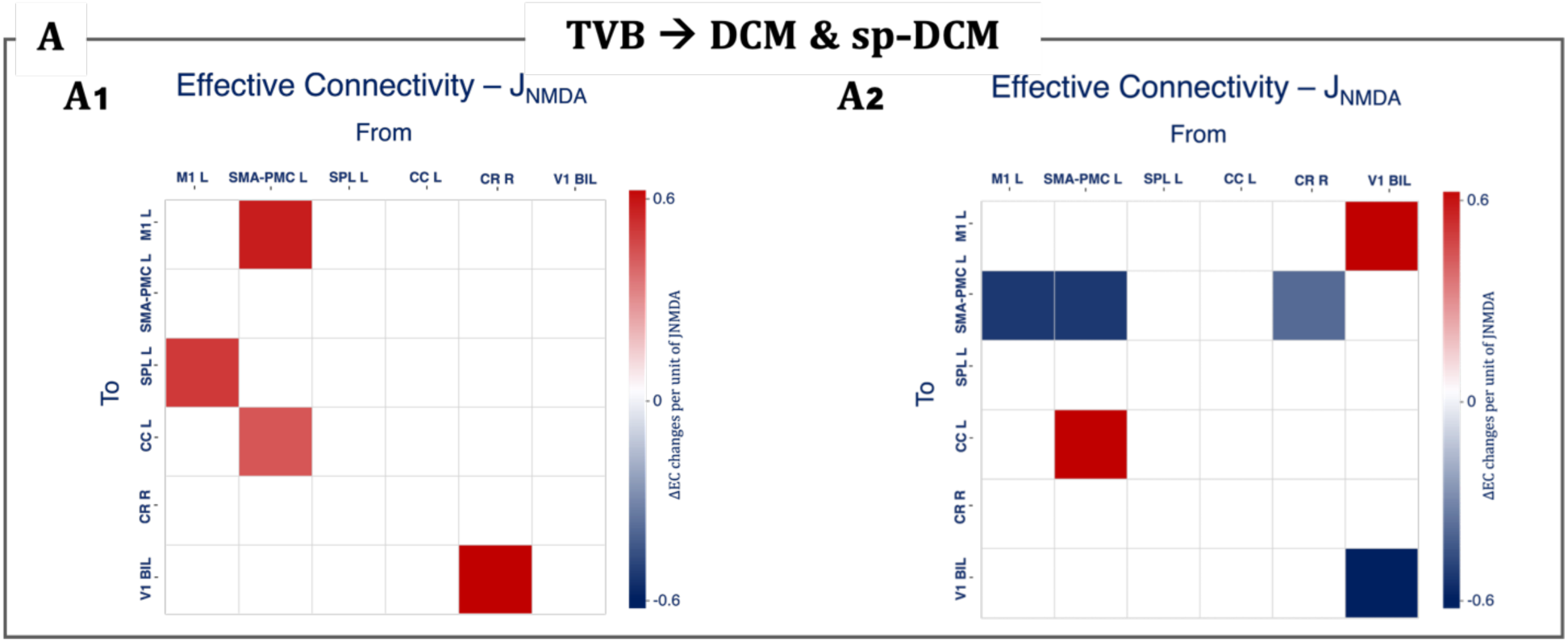
Post-hoc PEB analysis of J_NMDA_ - associated effective connectivity. PEB analysis with a posterior probability greater than 0.95 showed during task (A1), effective connectivity per unit of J_NMDA_ was increased from SMA-PMC to M1 and CC, from CR to V1, and from M1 to SPL. During rest (A2), higher J_NMDA_ was associated with reduced effective connectivity from sensorimotor regions (M1 and CR) to SMA-PMC and from, as well as with reduced SMA-PMC and V1 self-connectivity, and with increased effective connectivity from V1 to M1 and from SMA-PMC to CC. V1 = bilateral primary visual cortex, M1 = left primary motor cortex, SMA-PMC = left supplementary motor area and premotor cortex, CC = left cingulate cortex, SPL = left superior parietal lobule, CR = right cerebellar lobule VI.

Below, we describe these findings in detail.

### Network architecture

Figure 3A presents results identifying the best-fitting DCM functional model architecture for HV and pwMS. Between the 5 architectures tested for DCM, RFX-BMS identified the same winning model, S1.4, in both HV (A.1) and pwMS (A2) with the posterior probability exceeding 90%. In S1.4 V1 projects to regions involved in both motor planning and execution, including the CR, SMAPMC, and M1. The best-fitting model for the visuo-to-plan loop includes bidirectional connections between V1, CR, SMAPMC, and the CC, along with a self-inhibitory connection within the CR. M1 sits at the centre of this loop, receiving and sending bidirectional connections to V1, CR, and SMAPMC, and also connecting bidirectionally with visuomotor integration regions such as SPL (A3).

### Comparisons between HV and pwMS

DCM results of fixed effective connectivity (in Hz) for HV and pwMS are shown in Figure 3B. HV display stronger coupling strengths across most pathways than pwMS. The nature of the effective connectivity from SPL to M1 (identified by the asterisk * in Figure 3B) statistically significantly changes from being excitatory in HV to being inhibitory in pwMS (p=0.034).

sp-DCM results of fixed effective connectivity (in Hz) for HV and pwMS are shown in Figure 3C, estimated using the DCM winning network architecture (S1.4). Both groups exhibit non-symmetric and directed connectivity architectures. The nature of effective connectivity switches from excitatory to inhibitory or vice versa between HV and pwMS, with several statistically significant changes (identified by the asterisk * in Figure 3C) from M1 to SMAPMC (inhibitory in HV, excitatory in pwMS, p=0.008), from SPL to M1 (inhibitory in HV, excitatory in pwMS, p=0.010), from CR to V1 (inhibitory in HV, excitatory in pwMS, p=0.018) and from V1 to M1 (excitatory in HV, inhibitory in pwMS, p=0.045).

The relationship between GF and BOLD reflects task-dependent modulation of effective connectivity in the winning model (S1.4; S2), for both forward and backward pathways between HV (Figure 4A) and pwMS (Figure 4B). Winning-model architecture is identified following the stack procedure shown in Supplementary Figure 2, where each step’s winning model is identified with RFX-BMS (Supplementary Figure 3).

In HV, RFX-BMS identified the winning model for the forward pathway (from V1 to CR to SMAPMC to CC) with a probability >60% and the backward pathway (from CC to SMAPMC to CR to V1) with >70% (Figure 4A). Effective connectivity of the forward pathway showed positive modulatory effect with the GF-BOLD relationship, except for CR to SMAPMC, which showed negative modulation. Moreover, the forward pathway showed a linear GF-BOLD modulation from V1 to CR and from CR to SMAPMC, and nonlinear modulation from SMAPMC to CC, from V1 to SMAPMC and on the CR self-connection. All backward modulations showed a positive effective connectivity for an increased GF-BOLD correlation in HV, except for the V1 self-connection. The backward pathway presented linear modulation from CC to SMAPMC and the V1 self-connection, and nonlinear modulation from SMAPMC to CR and the CR self-connection. Figure 3B shows the forward and backward pathway models in MS, both selected by RFX-BMS with a probability >70%. In pwMS, the entire backward pathway showing a negative modulation on the effective connectivity, including the CR self-connection, and V1 self-connection showing a positive modulation. The same linear/nonlinear modulations are observed in pwMS as in HV, except from the connection from SMAPMC to CC, which is nonlinear in HV but linear in MS.

### Associations with clinical/behavioural measurements

To investigate how sensitive network dynamics are to disease severity, as measured by EDSS and reaction time, PEB analysis identified DCM and sp-DCM connections significantly associated with these clinical measures; a bidirectional stepwise regression was performed to examine how TVB parameter variance explained the same clinical/behavioural outcomes. All these results are reported in Figure 5.

PEB analysis identified DCM effective connectivity associations with EDSS (posterior probability > 0.95), (Figure 5A.1). In pwMS, higher EDSS (greater disability) was associated with reduced excitatory connectivity from SPL to M1, M1 to V1 and the V1 self-connection, as well as from CR to V1 and with increased effective connectivity from SMAPMC to M1 and to CR, from CR to SMAPMC, from V1 to M1, to SMAPMC and to CR. Effective connectivity associations with the reaction time (posterior probability > 0.95) are shown in Figure 5A.2. Longer reaction times were associated with reduced excitatory connections from M1 and CC to SMAPMC, from CR to V1 and the V1 self-connection, and with increased excitatory connections from CR to M1, CR self-connection, from V1 to M1 and to CR.

PEB analysis identified sp-DCM effective connectivity associations with EDSS (posterior probability > 0.95), (Figure 5B.1). Several connections within the visuomotor network showed significant associations with EDSS. Notably, there are several more connections in the resting-state condition, compared with task, that were associated with EDSS: higher EDSS scores (worse disability) were associated with reduced effective connectivity from SMAPMC to CR, From CC to SMAPMC and V1 self-connection, and with increased effective connectivity from M1 to SMAPMC, CR, V1 and M1 self-connection and from SMAPMC to M1 and CC, from CR and V1 to SMAPMC. Effective connectivity associations with reaction time (posterior probability > 0.95) are shown in Figure 5.B2. Longer reaction times were associated with reduced excitatory effective connectivity from M1 to CR, and from SMAPMC to M1 and with increased effective connectivity from M1 to SMAPMC, to SPL and to V1, from SMAPMC to CC and to CR, the CR self-connection, from V1 to CR and the V1 self-connection.

For the simulations of brain dynamics of the overall networks, all TVB parameters were normally distributed (p>0.05). Linear regression models were estimated with age, sex and TVB parameters (*G, J_NMDA_, J_i_, w_+_*) as independent variables and EDSS or reaction time as response variables. Between the four parameters examined, none demonstrated a statistically significant association with EDSS. The recurrent excitatory strength (*w_+_*) showed weak evidence for an association with EDSS (p = 0.07), although it did not meet the inclusion threshold of the final model. The excitatory glutamatergic coupling parameter (*J_NMDA_*) was a significant predictor of reaction time (β = -27.72, SE = 12.11, 95% CI [-53.8, -1.7], p = 0.034), explaining 21.6% of the variance (R² = 0.216, adjusted R² = 0.175), indicating that subjects with lower excitatory synaptic gain performed the task with longer delays, i.e. slower Figure 5.C).

A post-hoc analysis was performed to examine whether effective connectivity between pairs of regions was associated with the network-level J_NMDA_ parameter from TVB. During task, effective connectivity per unit of J_NMDA_ was increased from CR to V1, from SMA-PMC to M1 and CC and from M1 to SPL (Figure 6A.1). During rest, higher J_NMDA_ was associated with reduced effective connectivity from sensorimotor regions (M1 and CR) to SMA-PMC, as well as SMA-PMC and V1 self-connection, and with increased effective connectivity from V1 to M1 and from SMA-PMC and CC, reflecting a stronger interaction between the sensorimotor cortex and the cingulate cortex (Figure 6A.2).

## 4. Discussion

Our principal finding is that DCM, spDCM, and TVB can be brought together within a unified modelling framework that links effective connectivity to global-network dynamics. By linking connection-level estimates to emergent network dynamics, this approach allowed us to characterize differences between HV and pwMS in the visuomotor network and to explore the inferred excitatory-inhibitory balance underlying interregional interactions.

### Relationship between connection-level and network-level descriptions

A central observation is that robust differences between HV and pwMS emerged at the level of pairwise effective connectivity, while global TVB parameters did not show significant group differences. This apparent dissociation highlights an important methodological point: alterations in directed interactions between regions do not necessarily translate into measurable shifts in global network parameters, at least within the constraints of the present model and sample size.

From a modelling perspective, DCM and TVB provide complementary but non-equivalent descriptions of brain dynamics. DCM estimates context-dependent, directed coupling between predefined regions based on observed fMRI signals (Friston et al., 2003), while TVB simulates emergent network dynamics from structural connectivity and biophysically interpretable parameters (Ritter et al., 2013; Sanz-Leon et al., 2013). The present results suggest that local reconfigurations of effective connectivity may occur without consistent shifts in global dynamical regimes, or that such shifts are more difficult to detect given current model parameterisations. This is consistent with previous work showing that structural damage and functional reorganisation in MS can manifest heterogeneously across scales (Kennedy et al., 2024; Martí-Juan et al., 2023).

### Context-dependent modulation of effective connectivity

Across both groups, effective connectivity differed between task and resting-state conditions, confirming that interregional interactions are strongly context dependent. In HV, resting-state dynamics were characterised by predominantly inhibitory interactions, whereas task engagement was associated with a shift towards excitatory influences within the visuomotor network. This pattern is consistent with the interpretation of resting-state activity as a constrained regime that limits excessive signal propagation while maintaining readiness for task engagement.

In pwMS, this context-dependent modulation was altered. In particular, several connections showed reversals in the inferred sign (excitatory vs inhibitory) between groups, notably within cerebellar-cortical pathways. These findings extend previous fMRI studies demonstrating altered task-related responses and nonlinear scaling with motor demand in MS (Alahmadi et al., 2017, 2021) by suggesting that such differences may reflect changes in the organisation of directed interactions rather than solely in regional activation amplitudes.

### Cerebellar-cortical interactions as a locus of reconfiguration

The most consistent differences between groups involved cerebellar-cortical interactions, particularly the connection from cerebellar lobule VI (CR) to V1. During rest, this connection was inferred as inhibitory in HV but excitatory in pwMS, indicating a qualitative change in the direction of influence. Additional alterations in task-dependent modulations further implicated the cerebellum, including differences in the sign of feedback pathways and self-connections.

These findings are consistent with prior work highlighting the role of the cerebellum in coordinating visuomotor processing and adapting motor outputs to task demands (Diedrichsen et al., 2007; Lorenzi et al., 2025; Wolpert et al., 1998). In the context of MS, cerebellar involvement has been associated with both structural damage and functional reorganisation (Dogonowski et al., 2013; Savini et al., 2019). The present results extend these observations by suggesting that cerebellar contributions may be altered at the level of directed interactions, potentially influencing how sensory information is integrated into motor planning.

Previous studies have linked glutamatergic and GABAergic mechanisms to functional changes in MS within sensorimotor and parietal regions. Reduced glutamate has been associated with demyelination, while elevated GABA has been reported in sensorimotor areas, consistent with cortical reorganisation during disease progression (Bhattacharyya et al., 2013; Cawley et al., 2015; Macmillan et al., 2016; Nantes et al., 2017; Orije et al., 2015.) While DCM does not measure neurotransmitter concentrations directly, alterations in the sign and strength of effective connectivity may provide model-based evidence of such imbalance. The present findings are consistent with this picture, suggesting that disease-related shifts in excitatory and inhibitory coupling within the visuomotor network may reflect underlying neurochemical alterations, the temporal evolution and clinical relevance of which warrant further investigation.

### Associations with behaviour and clinical measures

Both DCM-derived effective connectivity and TVB parameters showed associations with behavioural performance and clinical disability. In particular, reaction time was related to the strength and sign of multiple connections, as well as to the NMDA-mediated excitatory gain parameter (J_NMDA_) in TVB. Participants with lower excitatory gain exhibited slower responses, suggesting that reduced effective integration within the network is associated with decreased behavioural efficiency.

Similarly, several connections were associated with EDSS, indicating that disease severity relates to distributed changes in network interactions rather than to single pathways. These findings are in line with previous studies reporting links between functional connectivity and clinical impairment in MS (Giorgio et al., 2010; Rocca et al., 2012), and support the view that network-level alterations contribute to clinical variability.

However, these associations should be interpreted with caution given the modest sample size and the number of parameters tested. The results therefore provide evidence of sensitivity rather than definitive biomarkers, and require validation in larger cohorts.

### Linking connection-level effective connectivity to network-level dynamics

A key insight from the combined DCM-TVB analysis is the relationship between global excitatory gain (J_NMDA_) and context-dependent reconfiguration of effective connectivity. J_NMDA_ values, obtained by optimising the excitatory synaptic gain at the network level, explained effective connectivity values across specific pathways during task, including from CR to V1 and projections from SMAPMC to M1 and CC. Within the constraints of the models used, this pattern suggests that greater baseline excitatory drive may require stronger directed coupling during task execution to sustain effective sensorimotor integration, particularly along motor planning (i.e. SMAPMC to M1), visuomotor integration and monitoring (SMAPMC to CC, M1 to SPL) pathways, as well as cerebellar engagement from early visual regions (CR to V1).

In contrast, during resting-state, J_NMDA_ was associated with a more differentiated pattern: a reduced interaction from distributed sensorimotor regions (M1 and CR) to SMAPMC, alongside increased coupling among visuomotor pairs (e.g., V1-M1, SMAPMC-CC). This suggests that, in the absence of task demands, network dynamics may favour intrinsic visuomotor coupling and internal evaluation processes, with relatively weaker integration of distributed sensorimotor inputs into motor planning and control hubs. The increased SMAPMC to CC interaction at rest may therefore reflect a greater coupling between action planning and evaluative systems, consistent with ongoing monitoring or readiness processes in the absence of overt behaviour.

Notably, cerebellar interactions showed no significant associations in the resting-state condition, in contrast to their engagement during task, consistent with the reduced need for online sensory feedback and error correction when no task is being performed. Together, these findings highlight that DCM-derived effective connectivity captures context-specific reweighting of directed interactions, while TVB parameters such as J_NMDA_ provide a more global constraint on network excitability. Their integration suggests that global synaptic gain does not directly determine network behaviour but modulates the extent to which directed interactions must be engaged to meet task demands, particularly in pathways supporting sensorimotor integration and performance monitoring.

### Study Considerations & Future Directions

Several limitations should be considered. First, the sample size is relatively small, which limits statistical power and the generalisability of the findings. Second, the analysis focused on a predefined visuomotor network, potentially excluding regions that may also contribute to disease-related alterations. Given that the dentate nucleus represents the principal cerebellar output channel projecting to the cortical areas via synapses in the thalamus and basal ganglia (Dum & Strick, 2003; Hoshi et al., 2005), future studies should investigate their involvement to better characterize the pathways mediating cerebro-cerebellar dysfunction in MS. Third, the interpretation of excitatory and inhibitory interactions relies on model-based inference and cannot be directly equated with synaptic or neurotransmitter-level processes. Finally, the integration between DCM and TVB remains mainly conceptual, apart from the final PEB analysis, and further methodological work is needed to establish formal links between these frameworks.

## 5. Conclusion

In summary, by combining DCM, sp-DCM, and TVB within a single conceptual framework, this study illustrates the potential of integrating complementary modelling approaches to characterise brain dynamics across scales. The results in the small HV and MS study suggest that connection-level and network-level descriptions capture different aspects of disease-related alterations, and that combining them may provide a more complete account than either approach alone. We show that MS is associated with alterations of effective connectivity that are context-dependent and that may imply a change in the excitatory/inhibitory nature of network dynamics, particularly involving cerebellar-cortical interactions. These changes are detectable at the level of pairwise directed interactions but do not necessarily manifest as global shifts in network parameters. At group level, integrating DCM and TVB can indeed link network-level excitability to context-dependent reconfiguration of directed interactions. This opens new avenue to study brain dynamics and neurological conditions.

## Acknowledgements

We would like to thank Amy Jolly for her support for fmriprep application.

