## Supplementary Materials for "A unified multiscale modelling framework to explore the brain excitatory-inhibitory balance: application to multiple sclerosis"

For clarity, the following summarizes the key preprocessing steps applied to anatomical and functional fMRI data using fMRIPrep 23.2.1

3DT1 images were put through a pipeline that included intensity non-uniformity correction, skull-stripping, and spatial normalization to the standard Montreal Neurological Institute template (MNI152NLin6Asym template) (Fonov et al., 2009). Then fMRI images were slice-time corrected, motion corrected, and co-registered to the corresponding 3DT1w data and normalized to the same MNI152NLin6Asym space. Outputs included confound time series of white matter and cerebrospinal fluid signal fluctuations that were used as regressors in the subsequent fMRI analysis. Resting state data was high-pass filtered with a 128-second cutoff to remove low-frequency drifts and the output MRI images, for both task and rest, were spatially smoothed with a 6 mm FWHM Gaussian kernel in SPM12.

### Supplementary Figures & Tables

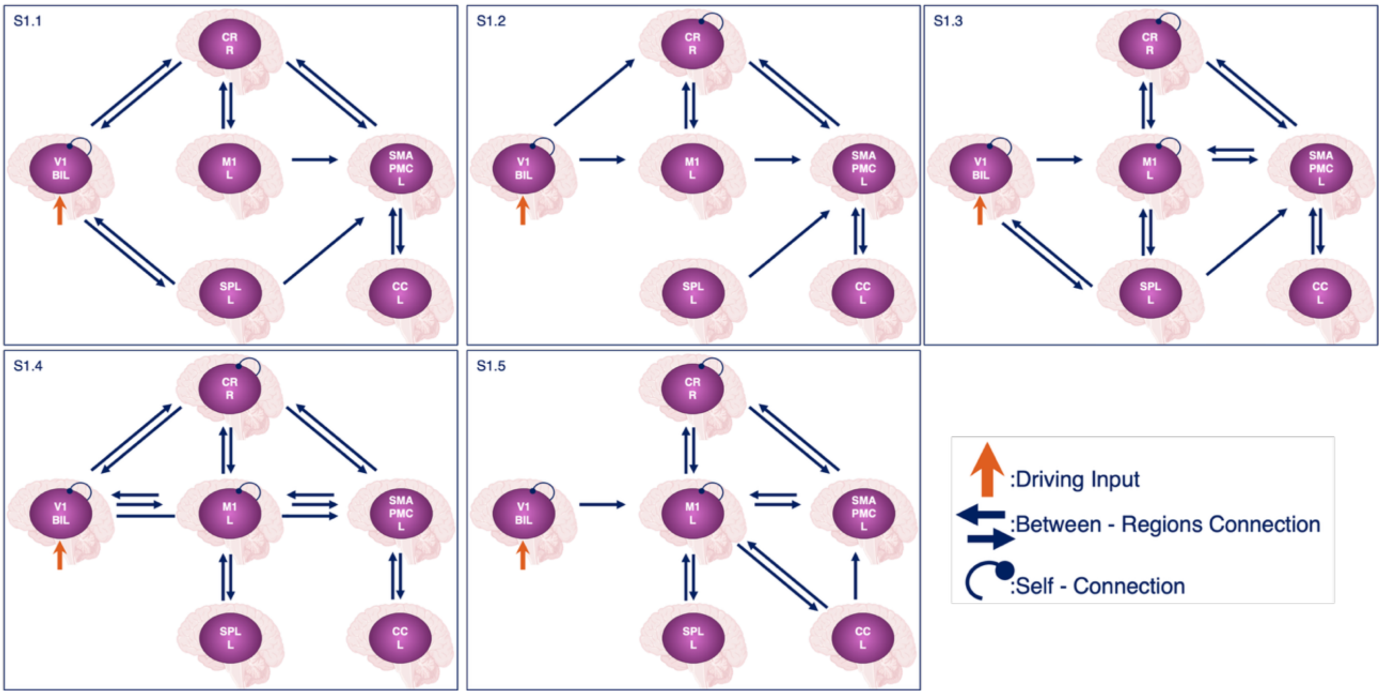


**Supplementary Figure 1. Visuomotor network models to assess effective connectivity.** The same five models obtained from a previously published visuomotor study by (Lorenzi et al., 2025b) are tested both for HV and pwMS, separately. The best model obtained with DCM for HV is also used in sp-DCM to investigate endogenous fluctuations in the resting-state condition. V1 = bilateral primary visual cortex, M1 = left primary motor cortex, SMAPMC = left supplementary motor area & premotor cortex, CC = left cingulate cortex, SPL = left superior parietal lobule, CR = right cerebellum.

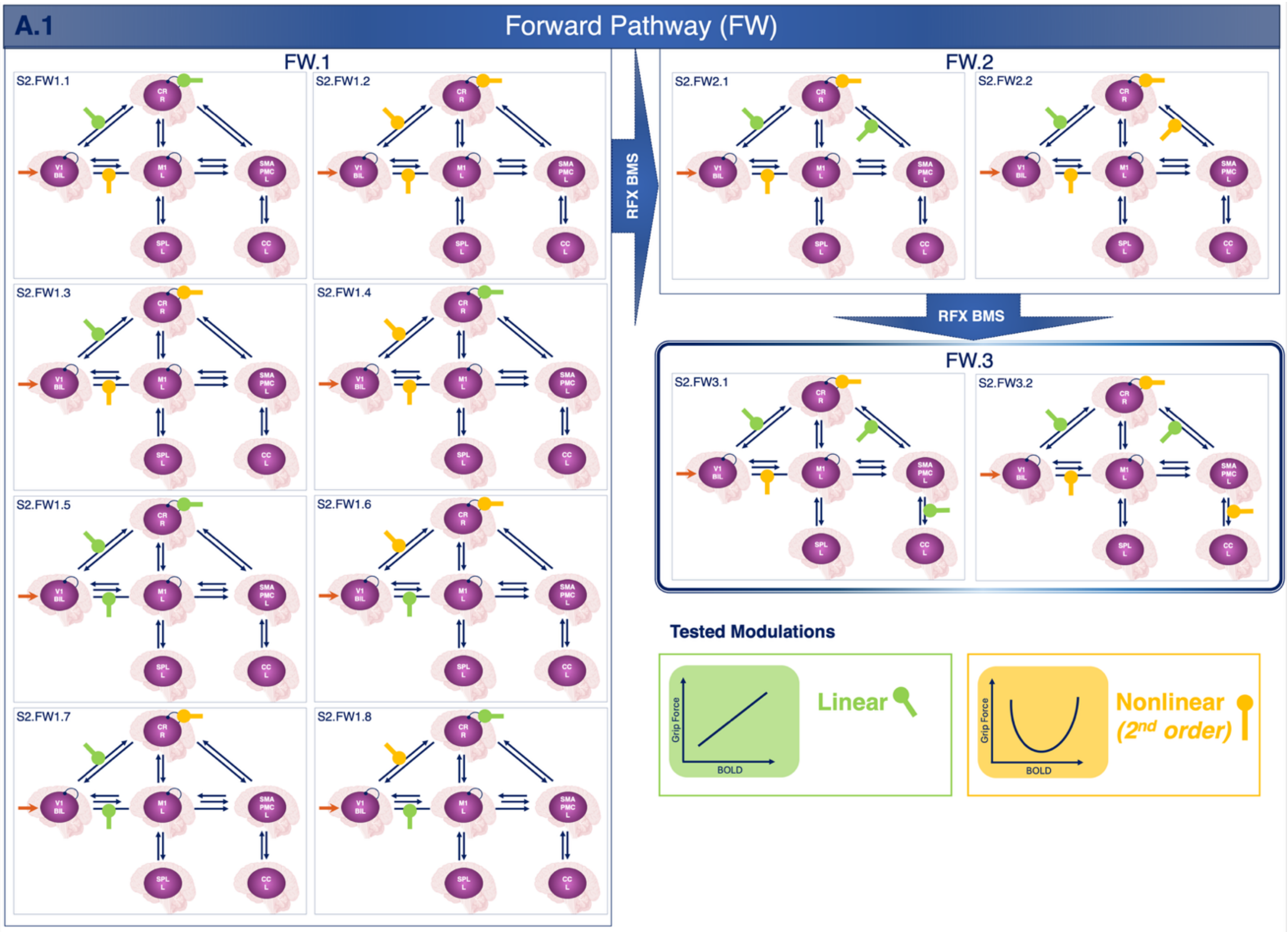
A.1
Forward Pathway (FW)
FW.1
S2.FW1.1
S2.FW1.2
S2.FW1.3
S2.FW1.4
S2.FW1.5
S2.FW1.6
S2.FW1.7
S2.FW1.8
RFX BMS
FW.2
S2.FW2.1
S2.FW2.2
RFX BMS
FW.3
S2.FW3.1
S2.FW3.2
Tested Modulations
Grip Force
BOLD
Linear
Grip Force
BOLD
Nonlinear
(2nd order)

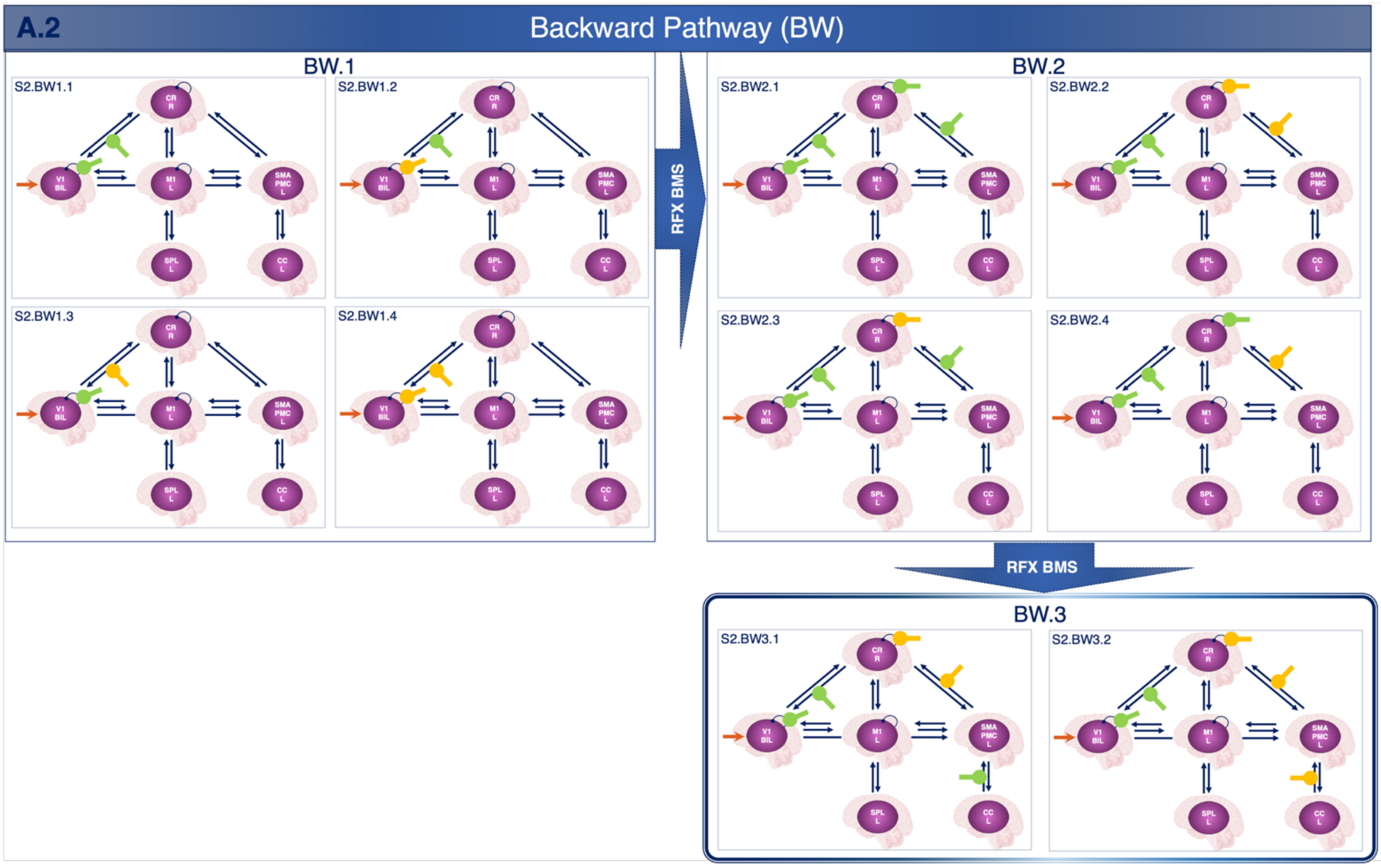
A.2
Backward Pathway (BW)
BW.1
S2.BW1.1
S2.BW1.2
S2.BW1.3
S2.BW1.4
RFX BMS
BW.2
S2.BW2.1
S2.BW2.2
S2.BW2.3
S2.BW2.4
RFX BMS
BW.3
S2.BW3.1
S2.BW3.2

**Supplementary Figure 2. Models to assess modulation of the fixed effective connectivity.** The winning model of the fixed effective connectivity (S1) is used in S2. Multiple configurations of GF-BOLD modulations are tested on the fixed effective connectivity for both pwMS and HV. The same strategy of linearity (1$^{st}$ order) and nonlinearity (2$^{nd}$ order) from (Lorenzi et al., 2025b) was applied. Forward (FW - A.1) and Backward (BW - A.2) pathways are defined focusing on the visuo-to-plan loop (V1-CR-SMAPMC-CC and back). A stack procedure is applied, grouping different modulations of the same connection and adding the winning modulation configuration before moving to the next group (starting from FW.1 and BW.1 towards FW.3 and BW.3 respectively). For both the FW and the BW pathways, the winning model of the most complex configuration was the overall winning model (FW3 and BW3 respectively). V1 = bilateral primary visual cortex, M1 = left primary motor cortex, SMAPMC = left supplementary motor area and premotor cortex, CC = left cingulate cortex, SPL = left superior parietal lobule, CR = right cerebellum.

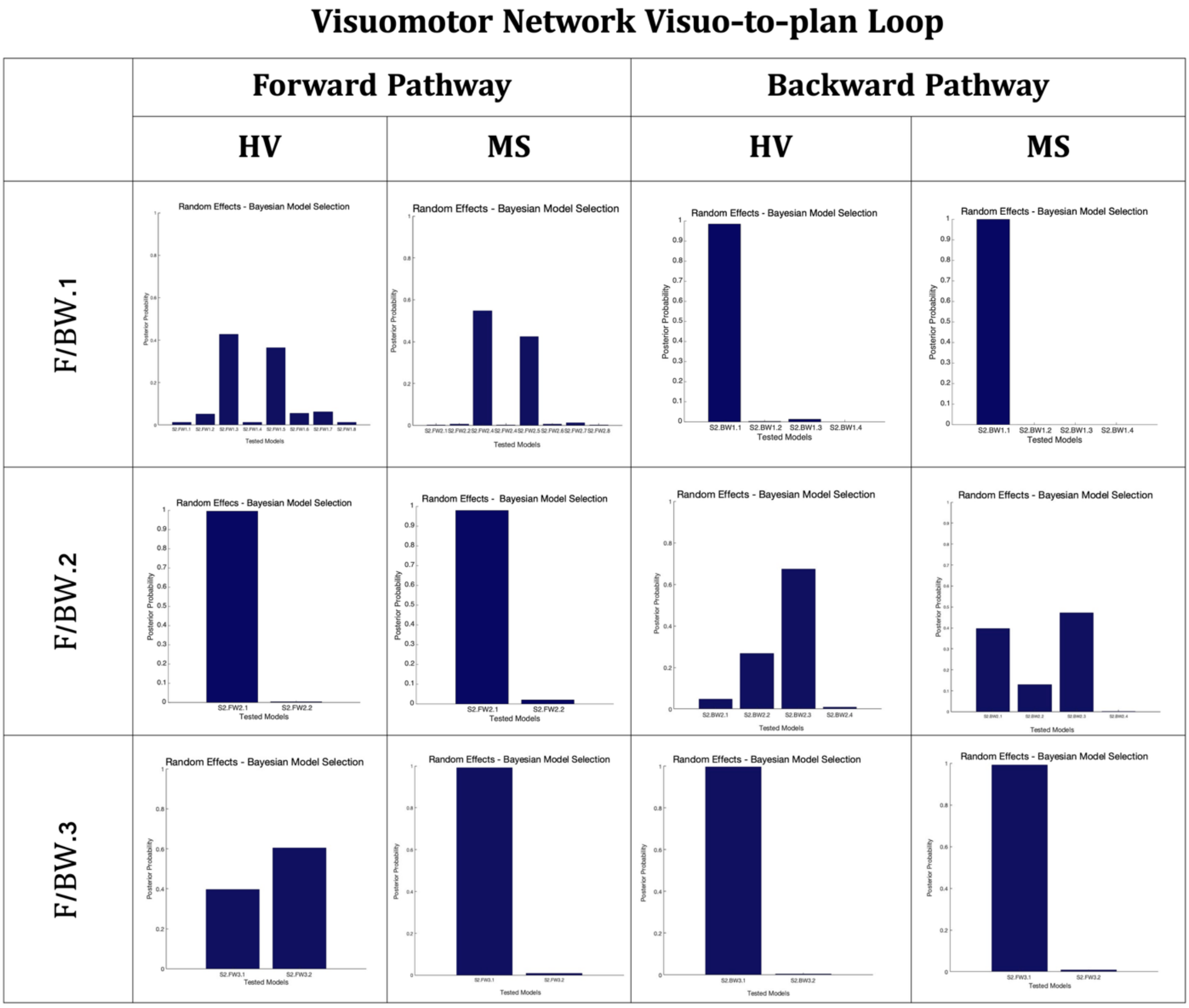


**Supplementary Figure 3. Each Forward and backward pathways assessed by RFX-BMS.** Random Effects Bayesian Model Selection results for each model family tested within the visuomotor "visuo-to-plan" loop. Each subpanel shows the posterior probability of the tested models for healthy volunteers (HV) and multiple sclerosis (MS) groups across forward and backward pathways. Model families (F/BW.1-3) were evaluated using Random Effects Analysis, and the winning models from each stage were subsequently entered into the stack procedure shown in Supplementary Figure 2 for the next level of model comparison.

**Table 1. MRI acquisition parameters for structural, task-based fMRI, resting-state fMRI, and diffusion imaging.**

| | Sequence | TR (ms) | TE (ms) | TI (ms) | Voxel size ($mm^3$) | Slices | Volumes/ Directions | FOV (mm) | Other parameters |
|---|---|---|---|---|---|---|---|---|---|
| PD/T2 (lesion) | Axial-oblique spin echo (dual-echo) | 3500 | 19 / 85 | - | 1 × 1 × 1 | 50 | - | 240 × 180 × 50 | Bicallosal alignment |
| T1-weighted | Sagittal-oblique 3D IR-FFE | 6.86 | 3.10 | 824 | 1 × 1 × 1 | 171 | - | 192 × 214 × 171 | Flip angle = 8° |
| Task fMRI | Axial-oblique T2* EPI | 2500 | 35 | - | 3 × 3 × 2.7 | 46 | 200 (+5 dummy) | 192 × 192 | Gap = 0.3 mm; SENSE = 2; descending; flip angle = 90° |
| Rs-fMRI | Axial-oblique T2* EPI | 2500 | 35 | - | 3 × 3 × 2.7 | 46 | 120 | 192 × 192 | Same parameters as task fMRI |
| Diffusion MRI | Axial-oblique HARDI (SE-EPI) | ~24* | 68 | - | 2 × 2 × 2 | - | 61 directions + 7 b0 | 112 × 112 × 72 | b = 1200 $s/mm^2$; cardiac-gated; flip angle = 90° |

* TR depended on heart rate due to cardiac gating.

**Supplementary Table 1. MRI Acquisition & fMRI Study Design.** A 3 Tesla Philips Achieva MRI scanner (Philips Healthcare, Best, The Netherlands) equipped with a 32-channel receive-only head coil was used for MRI acquisition on all the recruited participants. The imaging protocol included the following sequences; all aligned with the bicallosal line:

- Axial-oblique PD/T2-weighted spin echo sequence for lesion detection: dual-echo proton density (PD)/T2-weighted scans, echo time (TE1/TE2) = 19/85 ms, repetition time (TR) = 3500 ms, resolution = 1x1x1 $mm^3$, 50 slices, field of view (FOV) = 240x180x50 $mm^3$.
- Sagittal-oblique 3D T1-weighted (3DT1w) structural sequence: 3D inversion-recovery prepared gradient-echo (fast field echo) sequence with inversion time (TI) = 824 ms, TE = 3.10 ms, TR = 6.86 ms, voxel size = 1 × 1 × 1 $mm^3$, 171 slices, FOV = 192 x 214 x 171 $mm^3$, flip angle = 8°.
- Axial-oblique task fMRI data: BOLD sensitive T2*-weighted Echo Planar Imaging (EPI), TE = 35 ms, TR = 2500 ms, voxel size = 3 × 3 × 2.7 $mm^3$, inter-slice gap = 0.3 mm, SENSE factor = 2, 46 slices acquired in descending order, field of view (FOV) = 192 x 192 $mm^2$, number of volumes = 200, and 5 dummy scans, flip angle = 90°. The paradigm involved an event-related power grip task (Alahmadi et al., 2016), employing an fMRI-compatible squeeze ball with a visual cue, using the right (dominant) hand to squeeze a rubber ball with varying grip force levels. In detail, the task included 75 active trials, evenly but randomly distributed across five different levels of grip force: 20%, 30%, 40%, 50%, and 60% of each participant's maximum voluntary contraction. The actual timings of the task execution were recorded

for each subject and the delays between the task cue and the execution (i.e. the reaction times) were then calculated and z-score transformed.

- Axial-oblique rs-fMRI data: Geometrical and timing parameters were identical to the task fMRI acquisition, other than the number of volumes set to 120 as justified by (Van Dijk et al., 2010), showing that additional volumes did not significantly improve data quality in absence of the task.
- Axial-oblique DWI data: high angular resolution diffusion imaging (HARDI) were acquired with a cardiac-gated spin-echo (SE) EPI sequence with TE = 68 ms, TR= 24 (depending on heart rate) ms, voxel size = 2x2x2 $mm^3$, 61 isotropically distributed diffusion-weighted directions, b-value = 1200 $s/mm^2$, 7 b = 0 volumes, matrix size 112 x 112 x 72 $mm^3$, flip angle = 90°.
